# Bidirectional crusher gradient method for estimating the labeling efficiency of pseudo-continuous arterial spin labeling MRI in mice

**DOI:** 10.1101/2025.03.03.641196

**Authors:** Xiuli Yang, Yuguo Li, Adnan Bibic, Maria Guadalupe Mora Alvarez, Hanzhang Lu, Zhiliang Wei

## Abstract

Pseudo-continuous arterial spin labeling (pCASL) MRI is a widely used imaging technique for studying brain perfusion in health and disease due to its non-invasive and non-contrast nature. Accurate quantification of absolute perfusion values from pCASL signals requires the knowledge of labeling efficiency. However, to date, a reliable technique to measure pCASL labeling efficiency has not been available. In this study, we propose a method using bidirectional crusher gradients to modulate vascular signals in the azygos pericallosal artery (azPA) of the mouse brain, applied with and without pCASL labeling. The combination of corresponding signals allows the estimation of labeling efficiency. Upon systematic testing, optimal acquisition parameters included a labeling duration ≥ 1170 ms, a repetition time of 3 seconds, and an imaging slice thickness of 0.75 mm. In order to quantitatively estimate labeling efficiency, the bolus arrival time to azPA is required and found to be 218.7 ± 13.3 ms. Typical labeling efficiencies in mouse pCASL scans were 0.780 ± 0.048 (mean ± standard deviation). Furthermore, faster arterial flow induced by hypercapnia was found to increase pCASL labeling efficiency. Our method can improve the accuracy of pCASL quantification in mice, offering great potential for advancing its applications in pathophysiological studies.

## Introduction

Cerebral blood flow (CBF), which represents the amount of blood delivered to the brain per unit time, is a critical marker for studying various pathologies, including stroke, aging, Moyamoya, cardiac arrest, and Alzheimer’s disease.^1–5^ With a growing interest in monitoring CBF in both preclinical and clinical studies, there have been unprecedented efforts in developing novel imaging techniques for *in vivo* CBF assessment.^6,7^ Among these, arterial spin labeling (ASL) MRI has been extensively developed, rigorously optimized, and widely utilized.^8–10^ Its non-invasive, non-contrast nature makes it an indispensable tool in pathophysiological studies, particularly when longitudinal observation and subject comfort are priorities.^11^ ASL MRI relies on a kinetic model for accurate CBF quantification.^12^ Further refinement of the kinetic model to account for previously overlooked effects has enabled measurements beyond CBF, such as blood-brain barrier (BBB) function and cerebrospinal fluid (CSF) delivery rate.^13–15^ These advancements enhance the practical value of ASL MRI, establishing it as part of a comprehensive imaging toolkit for exploring various aspects of brain physiology.

There is a consensus to use pseudo-continuous ASL (pCASL) for perfusion imaging due to its pulse train design in the labeling module, which mitigates hardware requirements.^8^ Labeling efficiency, defined as the percentage of magnetization inverted by the labeling module, is essential for accurate modeling of pCASL signals.^12^ Numerical simulation was initially used to estimate labeling efficiency,^16,17^ but it soon became evident that experimental determination was necessary due to the intersubject differences, arterial location, and physiological variations. To address this, a method utilizing phase-contrast velocity MRI as a normalization factor was proposed.^18^ This approach maps the averaged ASL signal to the global CBF level under the assumption that water spins are fully extracted from the capillary bed to tissue – a well-accepted premise in ASL modeling^12^. However, the significantly lower water extraction fraction in mice (59.9 ± 3.2%^19^) compared to humans (95.5 ± 1.1%^20^) indicates that further technical development is needed before directly applying this normalization method for measuring labeling efficiency in mice. More recently, a method measuring blood magnetization at a downstream position near the labeling plane was proposed to estimate labeling efficiency in rats.^21^ By recording MRI signals in the complex mode, this method allows for the determination of magnetization polarity and, consequently, labeling efficiency after carefully addressing motion, pulsation, and eddy current artifacts etc.

In this study, we aim to develop a method for measuring the labeling efficiency of pCASL MRI in mice. Bidirectional crusher gradients were used to selectively retain and suppress ASL signals at the azygos pericallosal artery (azPA) in the midsagittal plane of the mouse brain, enabling quantification of inverted blood spins through pair-wise subtractions. Systematic optimizations were performed to establish an imaging protocol for measuring labeling efficiency, focusing on bolus arrival time to artery, dispersion effects from labeled-to-unlabeled blood exchanges during transition, repetition time, and slice thickness. The sensitivity of the proposed method to variations in labeling schemes was evaluated. Finally, the effect of hypercapnia on pCASL labeling efficiency was investigated.

## Material and methods

### Theory

Spin-echo echo-planar imaging (EPI) is widely used at ultrahigh magnetic fields to reduce image distortion caused by susceptibility inhomogeneity. In this study, we developed a method using bidirectional crusher gradients based on pCASL with spin-echo EPI acquisition (denoted as BIC-pCASL for descriptive convenience), as illustrated in Figure 1A. Crusher gradients flanking the refocusing pulse serve two functions: (a) dephasing unwanted signals caused by refocusing pulse imperfection and (b) selectively crushing blood signals flowing in the same direction as the crusher gradients. After combining control and labeled scans in pCASL, data acquisition is categorized into four scan types: Scan 1 corresponding to control scan with crusher gradients along the Z-axis (through-plane orientation), Scan 2 corresponding to labeled scan with crusher gradients along the Z-axis, Scan 3 corresponding to control scan with crusher gradients along the Y-axis (in-plane orientation), and Scan 4 corresponding to labeled scan with crusher gradients along the Y-axis (Figure 1B). The strengths and durations of the crusher gradients along different orientations are identical to ensure the same diffusion effect.

**Figure 1.**
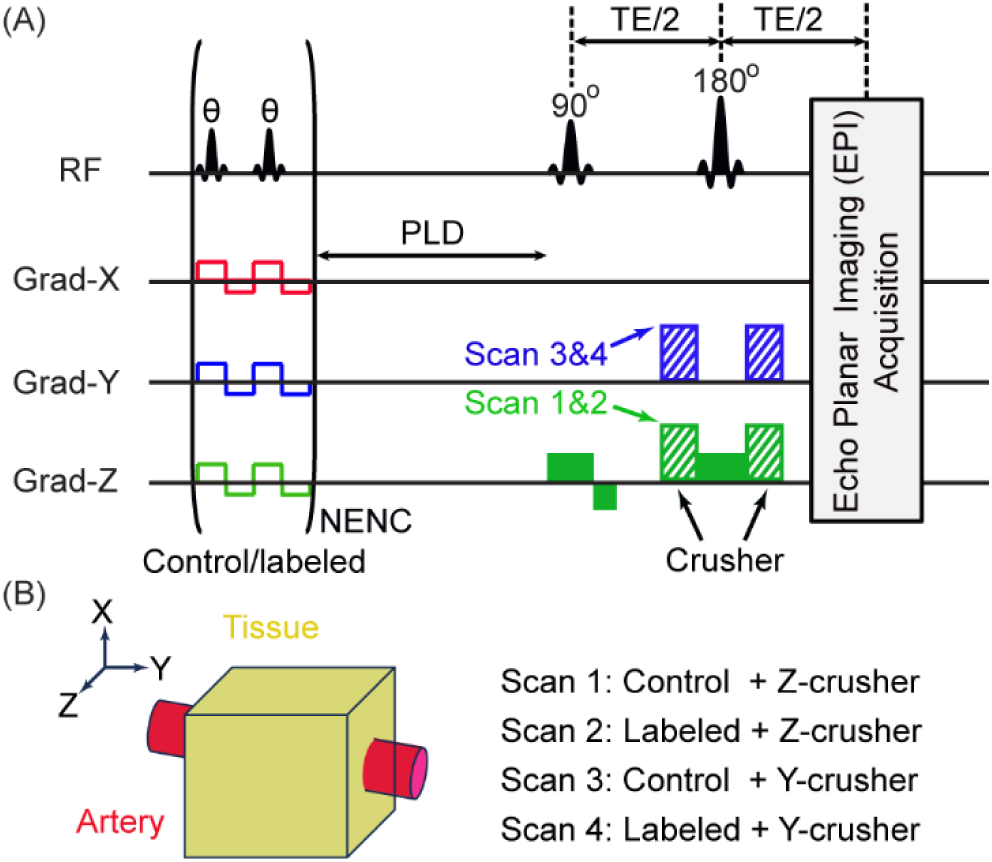
Schematic diagram of the BIC-pCASL method. (A) Pulse sequence of BIC-pCASL: NENC represents the number of labeling-pulse pairs (phases are 0° and 180° for the labeling pair in the control scan, and 0° and 0° for the labeled scan; when there is magnetic-field inhomogeneity, an additional phase will be applied). A spin-echo EPI is employed. Crusher gradients are applied along the Z axis in Scan 1& Scan 2 and along the Y axis in Scan 3 & Scan 4 to selectively suppress arterial signals. (B) Scan scheme. The imaging slice is positioned to include the artery in the X-Y plane. Four types of scans are conducted: Scan 1 corresponding to control scan with Z-axis crusher gradients, Scan 2 labeled scan with Z-axis crusher gradients, Scan 3 control scan with Y-axis crusher gradients, and Scan 4 labeled scan with Y-axis crusher gradients.

Considering an artery within the X-Y plane (Figure 1B), the MRI signal within an imaging voxel originates from three sources: tissue signal (𝑀_𝑡𝑖𝑠𝑠𝑢𝑒_), perfusion signal from capillary to tissue (𝑀_𝑝𝑒𝑟𝑓𝑢𝑠𝑖𝑜𝑛_), and vascular signal (𝑀_𝑣𝑒𝑠𝑠𝑒𝑙_). Let *α* denote the labeling efficiency and *β* the crusher efficiency (i.e., the percentage of magnetization eliminated by the crusher gradients), the MRI signals for different scans can be expressed as follows:

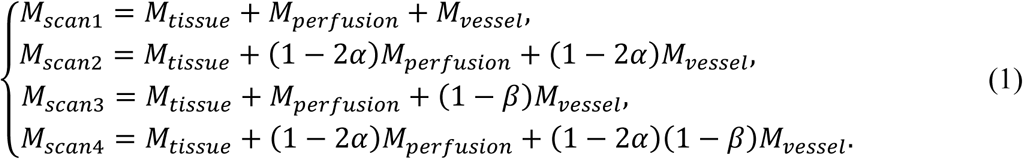

After performing pair-wise subtractions, the resulting difference signals can be expressed as follows:

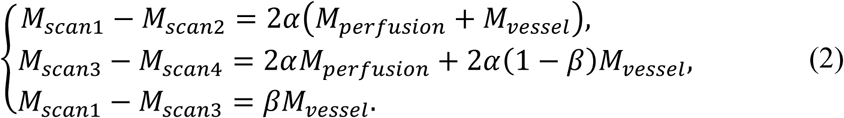

By further subtracting 𝑀_𝑠𝑐𝑎𝑛3_ − 𝑀_𝑠𝑐𝑎𝑛4_ from 𝑀_𝑠𝑐𝑎𝑛1_ − 𝑀_𝑠𝑐𝑎𝑛2_, Equation (2) turns into

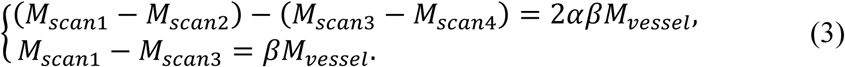

Thus, the labeling efficiency can be quantified as

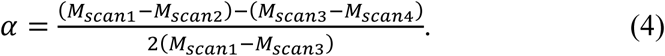

Note that the above derivations do not account for the *T*_1_ relaxation effect of blood spins during transit from the labeling location to the imaging voxel in the labeled scans. Assuming a bolus arrival time from the labeling location (neck region) to the imaging voxel (brain region) of 𝐵𝐴𝑇_𝑎𝑟𝑡𝑒𝑟𝑦_and an arterial *T*_1_ relaxation time of 𝑇_1,𝑏𝑙𝑜𝑜𝑑_, the labeling efficiency of pCASL MRI can be rewritten as

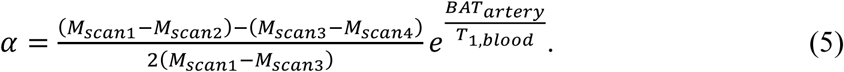

### MRI experiments

The experimental protocols for this study were approved by the Johns Hopkins Medical Institution Animal Care and Use Committee and conducted in accordance with the National Institutes of Health guidelines for the care and use of laboratory animals. Data reporting complied with the ARRIVE 2.0 guidelines. All procedures were carefully designed to minimize discomfort and stress to the animals. Mice were housed in a quiet environment under a 12-h light/dark cycle with ad libitum access to food and water. A total of 37 experimental sessions were conducted on a cohort of eight C57BL/6 mice (age: 37-41 weeks; body weight: 26-42 gram; 4 females, 4 males). Since MRI is a non-invasive technique, mice were used in multiple experimental sessions. For mice scanned on the same day, the experimental order was randomized according to a previously reported scheme.^22^

All MRI experiments were conducted on an 11.7T Bruker Biospec system (Bruker, Ettlingen, Germany) with a horizontal bore and an actively shielded pulse field gradient (maximum intensity of 0.74 T/m). Images were acquired using a 72-mm quadrature volume resonator as the transmitter and a four-element (2×2) phased-array coil as the receiver. Magnetic field homogeneity over the mouse brain was optimized using global shimming (up to the second order) based on a pre-acquired subject-specific field map. To minimize stress and motion, mice were anesthetized with inhalational isoflurane delivered via medical air (21% O_2_, 78% N_2_) at a flow rate of 0.75 L/min. Anesthesia was induced with 1.5-2.0% isoflurane for 15 minutes. At the 10^th^ minute of induction, the mouse was placed onto a temperature-controlled, water-heated animal bed and secured with a bite bar, ear pins, and a custom-built 3D-printed holder before entering the magnet. After induction, isoflurane concentration was reduced to 1.0% for maintenance during MRI scans and adjusted slightly (1.10-1.25%) if the respiration rate exceeded 120 breaths per minute. Respiration was observed throughout the experiment using an MR-compatible monitoring and gating system (SA Instruments, Stony Brook, USA). Experiments were terminated if respiration rates dropped below 50 breaths per minute for more than 2 minutes.

### Study 1: Determination of bolus arrival time to artery (N=8)

According to Equation (5), the 𝐵𝐴𝑇_𝑎𝑟𝑡𝑒𝑟𝑦_ value was required for estimating labeling efficiency and was the focus of Study 1. This study included four female and four male mice. A two-scan pCASL method, designed to minimize the influence of magnetic field inhomogeneity, was employed.^21^ First, a pre-scan was conducted to optimize the phases for labeling pulses in the control and labeled scans. Then, the pCASL scan was performed using the following parameters:^23^ repetition time (TR) / echo time (TE) = 3000 / 11.8 ms, field-of-view (FOV) = 15 mm × 15 mm, matrix size = 96 × 96, single slice covering the midsagittal plane, slice thickness = 1.0 mm, labeling pulse = 0.4 ms, inter-labeling-pulse delay = 1.0 ms, labeling duration = 300 ms, average number = 15, receiver bandwidth = 300 kHz, and scan duration = 3.0 min with 2-segment spin-echo EPI acquisition. To determine 𝐵𝐴𝑇_𝑎𝑟𝑡𝑒𝑟𝑦_, multiple post-labeling delays (PLDs) were used: 25, 50, 75, 100, 150, 200, 300, and 500 ms. Additionally, an M_0_ scan with a long TR of 20 seconds was acquired for normalization.

The processing of pCASL data followed established procedures.^22^ First, pair-wise subtraction between control and labeled images (i.e., 𝑀_𝑐𝑡𝑟_ − 𝑀_𝑙𝑏𝑙_) was performed to generate difference images, which were then normalized by corresponding *M*_0_ images to obtain perfusion-weighted images.^22^ A coarse region of interest (ROI) was manually drawn on the perfusion-weighted image to encompass the azPA, guided by a vasculature atlas.^24^ To ensure accurate arterial signal representation, the top ten voxels with the highest intensities within the ROI were automatically identified using MATLAB, and their average signal intensity was used as the arterial signal at azPA. The ROI for azPA was defined based on the averaged perfusion-weighted image across PLDs.

### Study 2: Optimization of labeling duration to ensure detection sensitivity (N=8)

Free diffusion between labeled and unlabeled blood spins can impact detection sensitivity, necessitating a sufficiently long labeling period. Therefore, Study 2 focused on optimizing labeling duration. This study included four female and four male mice. The pCASL scan, as described in Study 1, was performed with varying labeling durations (LDs) of 100, 200, 300, 400, 500, 600, and 1200 ms, while maintaining a constant PLD of 1 ms. In data processing, the ROI for azPA was defined based on the averaged perfusion-weighted image over the LDs.

### Study 3: Optimization of repetition time and slice thickness (N=8)

This study included four female and four male mice. To optimize TR, pCASL scans were conducted with two TR values (3 s and 6 s) while using an LD of 980 ms, a PLD of 20 ms, and an average number of 6. Additionally, to assess the effect of slice thickness, pCASL scans were performed with three slice thicknesses (0.50 mm, 0.75 mm, and 1.00 mm) while maintaining a TR of 3 seconds. In data processing, the ROI for azPA was delineated on the difference image and an ROI encompassing the cortex was drawn to compare tissue signals.

### Study 4: Sensitivity of labeling efficiency measurement to labeling schemes (N=7)

With the optimized parameters from Studies 1-3, the sensitivity of labeling efficiency to different labeling schemes was further examined. This study included four female and three male mice. Using the two-scan pCASL method, dynamic signals were measured as functions of phase offsets for both control and labeled scans. The phase offset combination yielding high control signals and low labeled signals was selected to maximize the difference signals, which are proportional to perfusion levels. Phase offsets of the labeled scans were changed to form different combinations and thereby different labeling schemes. For descriptive convenience, the labeling scheme producing 100% of the maximal difference signals was termed as optimal labeling (standard scheme in regular pCASL imaging); the labeling scheme producing approximately 50% of the maximal difference signals was termed as partial labeling; and the labeling scheme producing minimal difference signals was termed as minimal labeling (to be avoided in regular pCASL imaging). For each labeling scheme, a pCASL scan was performed to obtain whole-brain perfusion maps using the following parameters: TR/TE = 3000 / 11.8 ms, FOV = 15 mm × 15 mm, matrix size = 96 × 96, number of slice (axially) = 12, slice thickness = 0.75 mm, inter-slice gap = 0.25 mm, labeling pulse = 0.4 ms, inter-labeling-pulse delay = 1.0 ms, labeling duration = 1800 ms, PLD = 300 ms, average number = 25, receiver bandwidth = 300 kHz, and scan duration = 5.0 min with 2-segment spin-echo EPI acquisition. Additionally, BIC-pCASL was performed to measure labeling efficiency with the following settings: TR/TE = 3000/11.8 ms, slice thickness = 0.75 mm, labeling duration = 1200 ms, PLD = 100 ms, duration of crusher gradient = 2.0 ms, strength of crusher gradient = 0.148 T/m, readout orientation (X gradient) = ventral to dorsal, phase-encoding orientation (Y gradient) = rostral to caudal, slice selection orientation = left to right (Z gradient), average number = 8. All other BIC-pCASL parameters were identical to the pCASL scan.

Labeling efficiencies were quantified using Equation (5). The averaged 𝐵𝐴𝑇_𝑎𝑟𝑡𝑒𝑟𝑦_from Study 1 was used, and 𝑇_1,𝑏𝑙𝑜𝑜𝑑_was assumed to be 2813 ms based on literature reference.^25^ Perfusion-weighted images were co-registered and normalized to a mouse brain template^26^ to analyze averaged changing patterns.

### Study 5: Changes in labeling efficiency in response to hypercapnia challenge (N=6)

Hypercapnia is a widely used inhalational challenge for studying brain physiology. To facilitate future applications of pCASL in physiological studies, we investigated whether hypercapnia altered labeling efficiency in mice. This study included three female and three male mice. The pCASL and BIC-pCASL scans, as described in Study 4, were conducted under both normal air condition and pre-mixed gas containing 5% CO_2_ (21% O_2_ and 74% N_2_). The optimal labeling scheme was applied to maximize measurement sensitivity. For comparison, global cerebral blood flow (CBF) was measured using a previously reported phase-contrast (PC) MRI protocol.^27,28^ The key parameters for PC MRI were as follows:^29^ TR/TE = 15/3.2 ms, FOV = 15 mm × 15 mm, matrix size = 300 × 300, slice thickness = 0.5 mm, number of average = 4, dummy scan = 8, receiver bandwidth = 100 kHz, flip angle = 25°, encoding velocity = 20 cm/s (for left/right internal carotid arteries)/15 cm/s (for the basilar artery), partial Fourier acquisition factor = 0.7, and scan duration = 0.6 min per artery.

Absolute perfusion maps were quantified using the established equation:^8,21^ *CBF* = 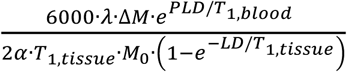, where *λ* represents the blood-brain partition coefficient (0.89 ml/g^30^), Δ𝑀 = (𝑀_𝑐𝑡𝑟_ − 𝑀_𝑙𝑏𝑙_), and 𝑇_1,𝑡𝑖𝑠𝑠𝑢𝑒_ denotes the *T*_1_ relaxation time of brain tissue. For the PC dataset, the artery of interest was manually delineated on the complex-difference image, which provided excellent contrast between the vessel and surrounding tissue.^27^ The resulting mask was applied to the velocity map, and arterial blood flow (in ml/min) was obtained by integrating the signal across arterial voxels. The total blood flow to the brain was then calculated by summing the flow values from the three major feeding arteries. To account for variations in brain size and derive unit-mass CBF values, the total blood flow was normalized by brain weight, which was determined as the product of brain volume and tissue density. The global CBF value was reported as milliliters per 100 grams of brain tissue per minute (ml/100g/min).^31^

### Data processing

All data processing was performed using custom-written MATLAB (MathWorks, Natick, MA) scripts and graphical user interface (GUI) tools.

### Statistical analyses

A two-way analysis of variance (ANOVA) was used to compare the Δ𝑀/𝑀_0_ signals between azPA and AIFA across PLDs in Study 1. A paired Student’s *t*-test was performed to compare: 𝐵𝐴𝑇_𝑎𝑟𝑡𝑒𝑟𝑦_ between azPA and AIFA (Study 1), cortex signals in control images between TRs (Study 3), azPA signals in difference images between TRs (Study 3), global CBF between gases (Study 5), inversion efficiencies between gases (Study 5), Δ𝑀/𝑀_0_signals between gases (Study 5), and mean flow velocities between gases (Study 5). A one-way ANOVA was conducted to compare: Δ𝑀/𝑀_0_signals across LDs (Study 2), cortex signals in control scans across slice thicknesses (Study 3), azPA signals in difference images across slice thicknesses (Study 3), Δ𝑀/𝑀_0_signals across labeling schemes (Study 4), and inversion efficiencies across labeling schemes (Study 4). Tukey’s Honest test was applied for post-hoc comparisons. Pearson correlation analysis was performed to assess the potential relationship between Δ𝑀/𝑀_0_ and labeling efficiency (Study 4). When reporting *t*-test results, *t*-statistics and degree of freedom (DF) were presented in the format *t* [DF]. For ANOVA results, DF between and within groups were reported in the format *F* [between-group DF, within-group DF]. When applicable, the 95% confidence interval (CI) was provided. All measurement values were presented as Mean ± Standard Deviation. A P value of <0.05 was considered statistically significant, with * indicating P<0.05, ** indicating P<0.01, and *** indicating P<0.001. Non-significant comparisons were denoted as n.s. for convenience.

## Results

### Study 1: Determination of bolus arrival time to artery

Figure 2 presents the results of PLD optimization. Pair-wise subtraction between control (Figure 2A) and labeled (Figure 2B) images produced difference images (Figure 2C) with clear contrast between arteries and surrounding tissues. Three arteries were identified in the midsagittal plane (Figure 2C) with reference to a vasculature atlas,^24^ including the anterior internal frontal artery (AIFA), azygos pericallosal artery (azPA), and basilar artery (BA). A signal built-up followed by a decay was observed in the Δ𝑀/𝑀_0_ signals at the group levels (Figures 2D and 2E). By fitting the Δ𝑀/𝑀_0_signals to Gaussian functions, the PLDs corresponding to peak Δ𝑀/𝑀_0_ signals were determined as 68.7 ± 13.3 ms for AIFA and 59.2 ± 19.1 for azPA (Figure 2D). Accounting for the labeling module duration, 𝐵𝐴𝑇_𝑎𝑟𝑡𝑒𝑟𝑦_ was estimated to be 218.7 ± 13.3 ms for AIFA and 209.2 ± 19.1 ms for azPA. These 𝐵𝐴𝑇_𝑎𝑟𝑡𝑒𝑟𝑦_values were not significantly different (*t* [7] = −1.23, P = 0.258). A two-way ANOVA further revealed that azPA exhibited significantly higher signals (P < 0.001). The BA, formed by the merging of the left and right vertebral arteries, is one of the three major feeding arteries, along with the left and right internal carotid arteries. Given that the internal carotid arteries contribute to 76.43 ± 1.35% of the total blood flow in mice across their major lifespan (Supplementary Figure S1), labeling efficiency estimation is more representative at a downstream branch of internal carotid arteries such as the azPA. Based on these results, azPA was selected for subsequent studies to improve signal-to-noise ratio (SNR) and representativeness.

**Figure 2.**
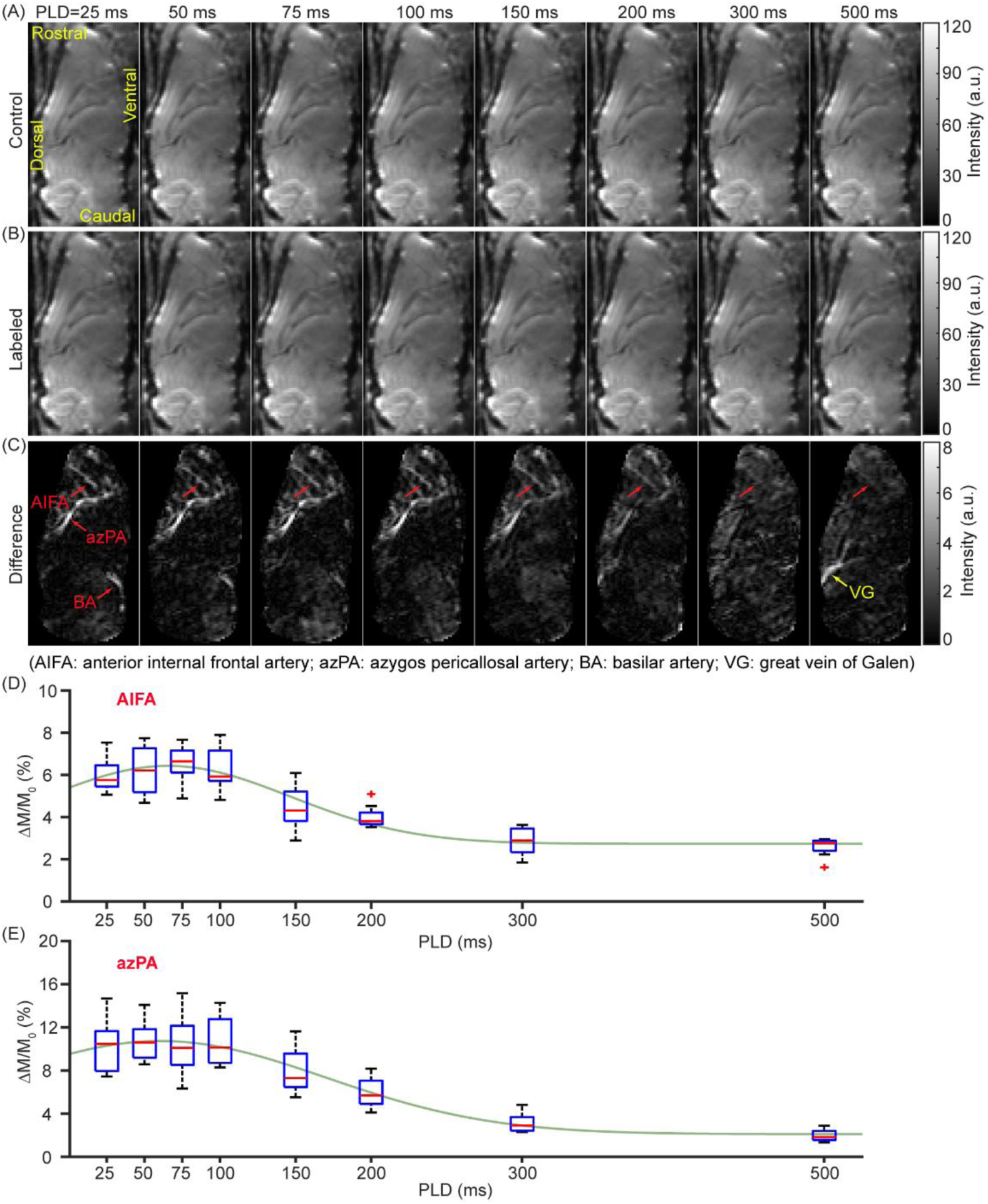
Determination of bolus arrival time to artery (N = 8). (A), (B), and (C) display the control, labeled, and difference images as functions of PLD. The anterior internal frontal artery (AIFA), azygos pericallosal artery (azPA), basilar artery (BA), and great vein of Galen (VG) were identified from the difference images. (D) and (E) illustrate the Δ𝑀/𝑀_0_signals as functions of PLD for AIPA and azPA, respectively. In the boxplot, the central red mark represents the median, while the top and down edges of the box correspond to the 25^th^ and 75^th^ percentiles. The whiskers extend to the minimal and maximal data points that are not considered outliers. The light green line represents the fitted gaussian function.

Bright signals were observed in the great vein of Galen (VG) at a PLD of 500 ms (Figure 2C), consistent with the previous report indicating a bolus arrival time to VG of 691.2 ms.^19^ Given the bolus arrival time to VG and an LD of 300 ms, a PLD of 541.2 ms was required. The 𝐵𝐴𝑇_𝑎𝑟𝑡𝑒𝑟𝑦_was significantly shorter than the bolus arrival time to VG (unpaired Student’s *t*-test, *t* [11] = −37.28, P < 0.001). Using the reported 𝑇_1,𝑏𝑙𝑜𝑜𝑑_ of 2813 ms,^25^ a scaling factor (i.e. 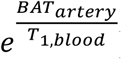) of 1.08 should be applied when estimating labeling efficiency from azPA.

### Study 2: Optimization of labeling duration to ensure detection sensitivity

Figure 3 presents the results of LD optimization. Control (Figure 3A) and labeled (Figure 3B) images of comparable quality were obtained across all tested LDs. In the difference images (Figure 3C), a progressive increase in MRI signals at the azPA was observed. ANOVA revealed a significant dependence of Δ𝑀/𝑀_0_ signals on LDs (Figure 3D, *F* [6, 49] = 40.01, P < 0.001). Since a labeled bolus undergoes free diffusion between labeled and unlabeled blood spins at both ends, shorter LDs were more susceptible to signal decay, as illustrated in Figure 3D. By fitting the Δ𝑀/𝑀_0_ signal into a Gaussian model, the full width at half height (FWHH) of dispersion was determined to be 326.2 ± 146.1 ms. The dispersion at azPA was significantly lower than that at the VG (806.4 ± 146.1 ms)^19^ (unpaired *t*-test, *t* [11] = −5.77, P < 0.001), primarily due to the significantly shorter bolus arrival time. Simulations based on the measured FWHH in individual mice predicted that an LD of at least 1170 ms would be required to achieve 99.9% of the maximal signal (Figure 3D).

**Figure 3.**
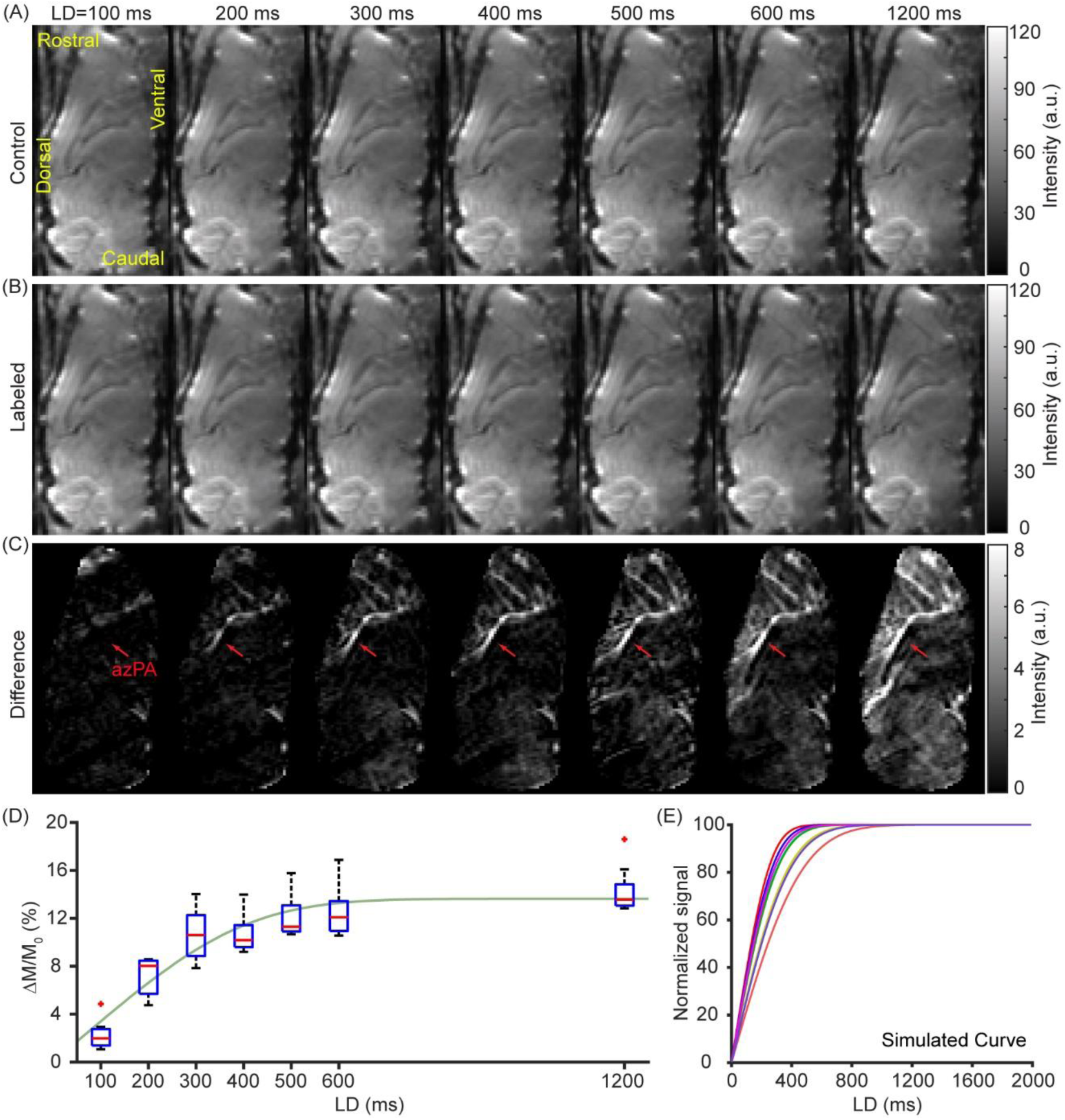
Determination of the full-width-at-half-height (FWHH) of dispersion effect (N = 8). (A), (B), and (C) show the control, labeled, and difference images as functions of LD. (D) illustrates the Δ𝑀/𝑀_0_ signal as a function of LD. (E) presents the simulated signals using the FWHH values obtained in (D).

### Study 3: Optimization of repetition time and slice thickness

The control and difference images primarily highlighted tissue and vascular signals, respectively. As shown in Figure 4A, the control image exhibited a larger change across TRs than the difference image. Two ROIs were selected to encompass the cortex and azPA regions (Figure 4B). Signal intensities of cortex were significantly higher in the control images at a TR of 6 s compared to 3 s (Figure 4C, *t* [7] = −4.25, P = 0.004), likely due to longer repetition time allowing for greater relaxational recovery. Difference signals at azPA remained unaffected by TR (Figure 4D, *t* [7] = 1.75, P = 0.122), indicating that a TR of 3 s was sufficient for flow-driven replacement of blood spins in azPA. Based on these findings, a TR of 3 s was selected for subsequent studies.

**Figure 4.**
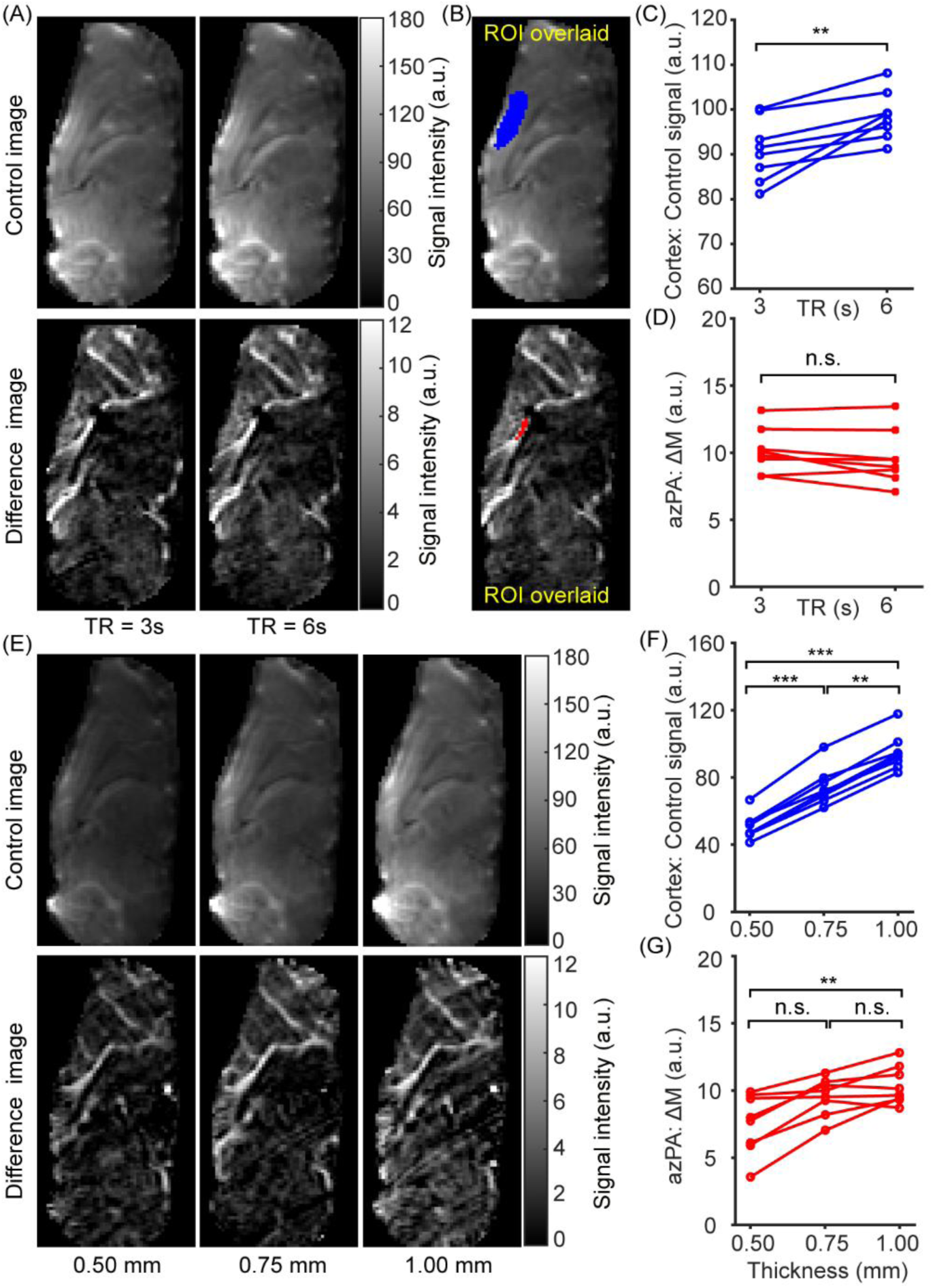
Optimization of repetition time and slice thickness (N = 8). (A) displays the control and difference images at two TRs. (B) shows ROIs for the cortex (blue) and azPA (red) overlaid on the control and difference images, respectively. (C) compares cortex signals in the control images across different TRs. (D) compares azPA signals in the difference images across different TRs. (E) presents the control and difference images across three slice thicknesses (0.50, 0.75, and 1.00 mm). (F) compares cortex signal in the control images across slice thicknesses. (G) compares azPA signal in the difference images across slice thicknesses.

As shown in Figure 4E, control images exhibited a clear increase in signal intensity as slice thickness increased. The azPA signal was successfully captured across all tested slice thicknesses (Figure 4E). At the group level, cortex signals in the control images varied significantly across slice thicknesses (*F* [2, 21] = 39.04, P < 0.0001). Tukey’s honest test further revealed significantly differences in cortex signals between 0.50 mm and 0.75 mm (95% CI = [-36.15, −11.03], P < 0.001), between 0.50 mm and 1.00 mm (95% CI = [-56.56, −31.44], P < 0.001), and between 0.75 mm and 1.00 mm (95% CI = [-32.97, −7.85], P = 0.001). These findings align with the fact that thicker slices encompass more brain tissue, leading to a greater number of protons contributing to MRI signals. Similarly, azPA signals showed a significant difference across slice thicknesses (*F* [2, 21] = 5.80, P = 0.010). Tukey’s honest test unraveled a significant difference between 0.50 mm and 1.00 mm (95% CI = [-5.01, −0.68], P = 0.009) but not between 0.75 mm and 1.00 mm (95% CI = [-3.02, 1.31], P = 0.589). There was a trend toward significance between the azPA signals of 0.50 mm and 0.75 mm (95% CI = [-4.16, 0.17], P = 0.074). Based on these results, a slice thickness of 0.75 mm was deemed sufficient to capture the azPA signal and was therefore used in subsequent experiments.

### Study 4: Sensitivity of labeling efficiency measurement to labeling schemes

Figure 5A illustrates an example of constructing labeling schemes by varying the phase offsets for labeled scans. The averaged pCASL signals (Δ𝑀/𝑀_0_) differed significantly across the labeling schemes (*F* [2, 18] = 50.03, P <0.001) (Figure 5B). Tukey’s honest test further revealed significant differences in pCASL signals between the optimal and partial labeling (95% CI = [0.79, 1.95], P < 0.001), between the optimal and minimal labeling (95% CI = [1.68, 2.84], P < 0.001), and between the partial and minimal labeling (95% CI = [0.31, 1.47], P = 0.003). Labeling efficiencies also exhibited a significant group-level difference (*F* [2, 18] = 97.84, P < 0.001) (Figure 5C). According to Tukey’s honest test, labeling efficiency significantly differed between the optimal and partial labeling (95% CI = [0.27, 0.51], P < 0.001), between the optimal and minimal labeling (95% CI = [0.54, 0.78], P < 0.001), and between the partial and minimal labeling (estimate = 0.26, 95% CI = [0.14, 0.38], P < 0.001) (Figure 5C). Additionally, a strong correlation was observed between Δ𝑀/𝑀_0_ and labeling efficiency (R^2^ = 0.860, P < 0.001).

**Figure 5.**
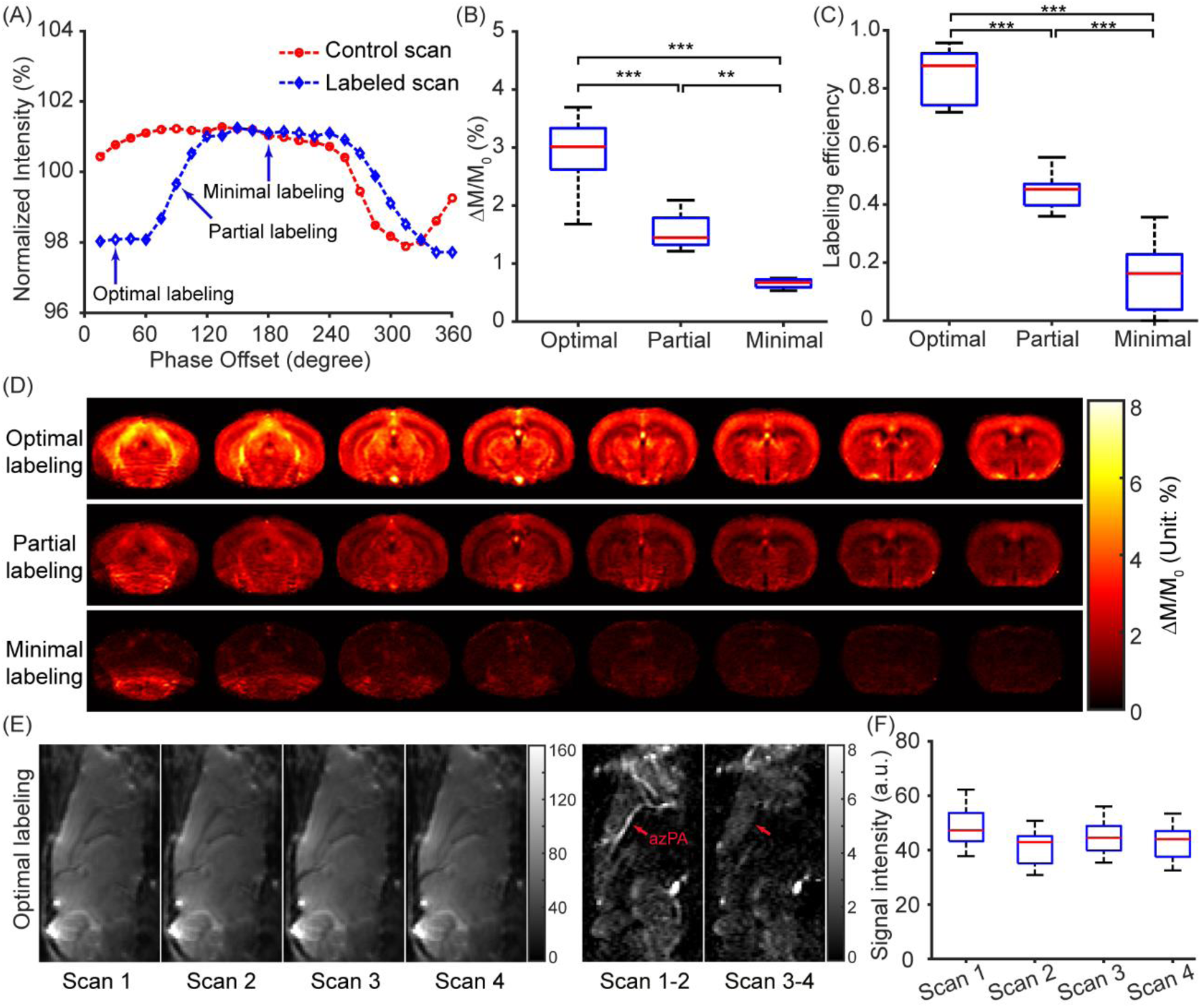
Sensitivity of labeling-efficiency measurements to labeling schemes (N = 7). (A) shows labeling schemes obtained by varying the phase offsets of labeled scans. Three labeling schemes were compared: optimal, partial, and minimal labeling. (B) and (C) compare Δ𝑀/𝑀_0_and labeling efficiency among the optimal, partial, and minimal labeling schemes, respectively. (D) shows Δ𝑀/𝑀_0_ images under the optimal, partial, and minimal labeling schemes. (E) presents an exemplary dataset of BIC-pCASL. Pair-wise subtraction highlights the retention (Scan 1 – Scan 2) and suppression (Scan 3 – Scan 4) of azPA signals under crusher gradients applied along different axes. (F) illustrates signal intensities at the azPA across different scan types.

A more intuitive observation on the influence of labeling schemes is presented by the averaged perfusion-weighted images in Figure 5D, showing a noticeable signal decay from optimal labeling to partial labeling, and further to minimal labeling. An exemplary dataset for BIC-pCASL is displayed in Figure 5E, where difference images obtained via pairwise subtraction illustrated selective vascular suppression under different crusher gradients. The azPA signals were preserved under crusher gradients applied along the through-plane orientation (Scan 1 – Scan 2, Figure 5E) but were suppressed under crusher gradients applied along the in-plane orientation (Scan 3 – Scan 4, Figure 5E). The original signal intensities of the azPA across different scans are illustrated in Figure 5F. There was a significant difference between the (Scan 1 – Scan 2) and (Scan 3 – Scan 4) signals (*t* [6] = 7.03, P < 0.001), which aligns with theoretical expectations. Moreover, there was no significant difference in signal intensities across the four scan types (*F* [3, 24] = 1.58, P = 0.219), indicating that MR signals in the azPA ROI primarily originated from sources insensitive to crusher gradients, such as tissue. The inverted blood spins were insufficient to alter the polarity of MR signals at the voxels, thereby avoiding potential misinterpretation of negative MR signals due to the use of magnitude mode in data acquisition.

These results highlight the importance of achieving high inversion efficiencies to enhance the sensitivity of perfusion maps. Additionally, labeling efficiency measured by BIC-pCASL proves to be highly responsive to variations in labeling conditions.

### Study 5: Changes in labeling efficiency in response to hypercapnia challenge

As illustrated in Figure 6A, regional perfusion maps demonstrated increased CBF across various brain regions under hypercapnia. PC MRI confirmed a significantly rise in global CBF (*t* [5] = −5.72, P = 0.002, Figure 6B), validating the vasodilatory effect of CO_2_. Labeling efficiencies significantly differed between medical air (0.780 ± 0.048) and 5% CO_2_ gas (0.845 ± 0.078) (*t* [5] = −3.05, P = 0.029, Figure 6C). In line with the increase in labeling efficiencies, Δ𝑀/𝑀_0_ was also significantly higher under hypercapnia (*t* [5] = −3.87, P = 0.012, Figure 6D). Concurrently, the mean flow velocities across major feeding arteries increased significantly during hypercapnia (*t* [5] = −2.94, P = 0.032, Figure 6E). Bloch simulations indicated that flowing velocity modulates adiabatic inversion in pCASL labeling, as evidenced by varying inversion levels at flowing velocities of 5, 10, and 15 cm/s (Figure 6F). Further simulations using the same parameters as the actual experiments, conducted over a velocity range of 0.5–30 cm/s, demonstrated a velocity-dependent pattern in labeling efficiency. Below 16.7 cm/s, labeling efficiency positively correlated with flowing velocity (R^2^ = 0.662, P < 0.001), whereas, beyond this threshold, a negative correlation emerged (R^2^ = 0.964, P < 0.001). The mean flow velocities observed under normocapnia and hypercapnia fell within the 0.5 – 16.7 cm/s range, corresponding to higher labeling efficiency at increased flowing velocity. Our experimental findings were consistent with simulation results. In summary, hypercapnia enhances the labeling efficiency of pCASL MRI in mice by accelerating arterial blood flow.

**Figure 6.**
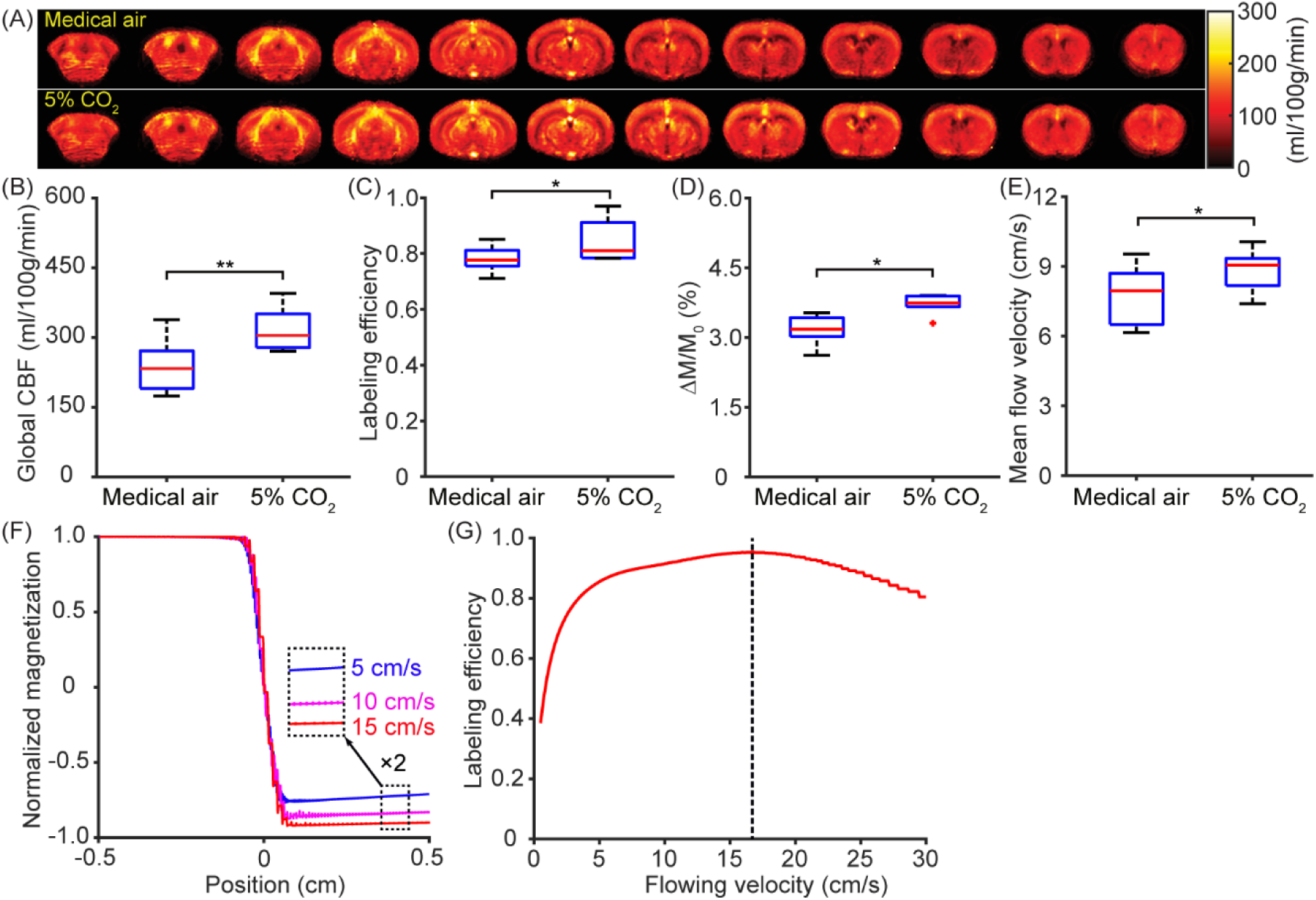
Influence of hypercapnia challenge on the labeling efficiency (N = 6). (A) shows regional perfusion maps under medical air and 5% CO2 gas. (B), (C), (D), and (E) show the comparisons of global CBF, labeling efficiencies, Δ𝑀/𝑀_0_, and mean flow velocities between medical air and 5% CO2 gas, respectively. (F) illustrates the flow-driven adiabatic inversion processes for blood spins at velocities of 5, 10, and 15 cm/s. (G) depicts the dependence of labeling efficiency on flowing velocity.

## Discussion

In this study, we introduced a method utilizing bidirectional crusher gradients to assess the labeling efficiency of pCASL MRI in mice, termed BIC-pCASL. Systematic optimization of key parameters was conducted to establish an optimized imaging protocol for BIC-pCASL. The labeling efficiency measured by BIC-pCASL demonstrated high sensitivity to variations in the labeling scheme. Additionally, hypercapnia was shown to enhance labeling efficiency by increasing arterial blood flow velocity.

CBF is a potential biomarker for monitoring vascular pathology, assessing therapeutic efficacy, and evaluating physiological perturbations following drug administration.^32–35^ Evaluating CBF in diseases characterized by vascular dysfunction can aid in both diagnosis and prognosis.^36,37^ For example, perfusion imaging helps delineate hypoperfusion boundaries in ischemic stroke, facilitating the identification of the core region and ischemic penumbra.^38^ Additionally, early-stage CBF recovery after cardiac arrest serves as a predictor of neurological outcomes.^33^ Due to neurovascular coupling, CBF alterations occur not only in vascular pathologies but also in metabolic disorders such as Alzheimer’s disease.^39,40^ Furthermore, CBF can be integrated into multiparametric studies alongside other imaging and physiological techniques to comprehensively characterize pathophysiological changes.^14,41–43^ Specially, combining vascular physiology with pathological assessments^44,45^ enhances our understanding of the full spectrum of pathophysiological changes.^46–48^

ASL MRI is a state-of-the-art, non-contrast technique for imaging perfusion. With its growing application in both preclinical and clinical studies, significant efforts have been made to enhance its quantification accuracy.^8,49–51^ Accurate measurement of labeling efficiency is particularly crucial for intersubject comparisons, where absolute perfusion values (unit: ml/100g/min) are desired. This is especially relevant in case such as using hippocampal CBF as a diagnostic aid for Alzheimer’s disease.^52^

The simulation method examines magnetization behavior under specific experimental parameters using the Bloch equations. It calculates the optimal labeling efficiency achievable for a given parameter combination. However, accounting for all experimental imperfections, such as field inhomogeneity and intersubject heterogeneity, remains challenging. For instance, precise positioning of the labeling slice is difficult because not all feeding arteries run parallel at the labeling position, and vascular tortuosity may be present within the labeling plane. Given these subject-specific variations, experimental determination of labeling efficiency for pCASL scans is preferred.^18^ Our study extends the previous experimental methods^18,21^ to estimate the labeling efficiency of pCASL MRI.

The normalization method, which references global CBF obtained via PC MRI, estimates labeling efficiency by assuming that the averaged ASL perfusion equals PC-based global CBF. In humans, the water extraction fraction from capillaries to tissue is 95.5 ± 1.1%^20^, supporting the key assumption required for normalization. Note that some discrepancies between pCASL and PC MRI results have been reported in human, with PC MRI consistently showing significantly higher CBF than pCASL MRI.^53^ These discrepancies arise from different systematic errors associated with PC and pCASL MRI. PC MRI tends to overestimate CBF due to imperfect slice positioning,^27^ whereas pCASL MRI often underestimates CBF due to overestimated labeling efficiency. Factors such as imperfect positioning of labeling plane, magnetic-field inhomogeneity, and vascular tortuosity etc. can reduce labeling efficiency, leading to an assumed or simulated labeling efficiency that is higher than the actual labeling efficiency. According to the equation 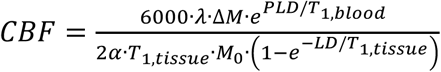, CBF is underestimated when labeling efficiency is overestimated. Given these opposing systematic errors, the observation that PC MRI yields significantly higher CBF than pCASL MRI is expected. In mice, the water extraction fraction is significantly lower, at 59.9 ± 3.2%^19^, meaning that only about half of the blood water is extracted by brain tissue. This reduction is primarily due to the use of isoflurane, a potent vasodilative anesthetic that minimizes motion and stress but accelerates blood flow, reducing the time available for water extraction by tissue.^19^ In human pCASL data, normalized regional maps reflect perfusion after water spins have been extracted from capillaries to tissue. In contrast, applying the normalization method to mouse pCASL data, the obtained results represent regional blood supply before tissue extraction. A recent study examining the pro-aging effects of excessive PDGF-BB suggests that the normalization method does not alter the findings related to vascular dysfunction.^54^ In summary, water extraction fraction must be carefully considered when implementing the normalization method, particularly in physiological conditions with reduced water extraction, such as in patients with sickle cell disease.^55^

A more recent approach for evaluating labeling efficiency involves measuring blood signals in the complex mode at a downstream position close to the labeling plane.^21^ In this method, magnetization polarity is reflected by the angle of the complex data. It was noted by the developer that unbalanced contributions between feeding carotid arteries could compromise quantification accuracy. In contrast, the BIC-pCASL method focuses on a distant artery (i.e., azPA) located farther from the labeling position. This design reduces vulnerability to imbalances across feeding arteries, as well as pulsation and respiration-related motion artifacts. Our proposed method serves as a valuable complement to existing techniques, enhancing the reliability of labeling efficiency evaluation in pCASL MRI across various applications.

In this study, we focused on the azPA because it provides better detection sensitivity than the AIFA at the midsagittal plane of the mouse brain. However, if a mouse exhibits an abnormal vascular trajectory that prevents the observation of the azPA under a midsagittal imaging slice, the AIFA can be analyzed as an alternative for estimating labeling efficiency. Since the bolus arrival time to the AIFA is similar to that of the azPA, the scaling factor 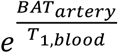 remains 1.08 for AIFA.

Selective crushing of arterial MR signals in difference scans forms the technical foundation of BIC-pCASL. In principle, higher crushing efficiency results in larger differences between the (Scan 1 – Scan 2) and (Scan 3 – Scan 4) signals, thereby enhancing detection sensitivity. However, complete crushing of arterial signals in the azPA is not required for implementing BIC-pCASL, as labeling efficiency is measured in a relative percentage manner according to Equation (4). On the other hand, changes in magnetic field strengths will not compromise the technical foundation of BIC-pCASL. Bolus arrival time to azPA, exchange between labeled and unlabeled blood spins, replacement of blood spins in azPA, and vascular dimension of azPA are independent of magnetic-field strength. Therefore, generalizing BIC-pCASL to field strengths other than 11.7T should be feasible.

Monte Carlo simulations with 2,000 iterations were conducted to evaluate statistical power. Based on the means and standard deviations of the collected MRI data, the statistical power was calculated for various comparisons: 0.99 for detecting differences between AIFA and azPA signals (Study 1), 0.79 for detecting differences in azPA signals across slice thicknesses in (Study 3), 0.99 for detecting differences in Δ𝑀/𝑀_0_signals across labeling schemes (Study 4), 0.99 for detecting differences in labeling efficiency across labeling schemes (Study 4), 0.73 for detecting differences in labeling efficiency across gas conditions (Study 5), and 0.70 for detecting differences in Δ𝑀/𝑀_0_ signals across gas conditions (Study 5) with the given sample sizes. These results indicate that the findings in this study were supported by sufficient statistical power.

Results from this study should be interpreted considering certain limitations. While the radiofrequency power deposition of pCASL is lower than that of continuous ASL, it remains higher than that of pulsed ASL, necessitating careful optimization of parameters such as labeling duration and inter-labeling-pulse delay etc. Incorporating the BIC-pCASL scan extends the total experimental duration. In applications requiring rapid imaging, such as acute-stage reperfusion after the return of spontaneous circulation in cardiac arrest, the need for an additional scan may reduce the overall temporal resolution of perfusion imaging. Recent advancements in signal processing, particularly artificial intelligence algorithms, have been demonstrated to enhance MRI performance^56–58^ and could be explored to improve the temporal resolution of pCASL MRI in these time-sensitive applications.

In summary, we introduced a method utilizing bidirectional crusher gradients to estimate the labeling efficiency of pCASL MRI in mice by tracking blood spins at the arterial side. Through systematic optimization and validation studies, we established the methodology and determined a typical labeling efficiency of 0.780 ± 0.048 for pCASL scans in mice. Hypercapnia was found to enhance labeling efficiency by accelerating blood flows in arteries. This proposed method strengthens the accuracy of pCASL-based perfusion quantification in mice and hold promise for advancing the application of pCASL in future pathophysiological studies using animal models.

## Acknowledgements

This work was supported by the National Institutes of Health (NIH) under R01 AG081932, P41 EB031771.

## Authors’ contributions

X.Y.: Methodology, Investigation, Formal analysis, Visualization, Data Curation, Writing – Original Draft, and Writing -Review & Editing. Y.L.: Investigation and Writing -Review & Editing. A.B.: Investigation and Writing -Review & Editing. M.M.A.: Formal analysis and Writing -Review & Editing. H.L.: Formal analysis, Resources, Funding acquisition, and Writing -Review & Editing. Z.W.: Conceptualization, Methodology, Investigation, Formal analysis, Visualization, Data Curation, Resources, Funding acquisition, and Writing - Review & Editing.

## Declaration of conflicting interest

The author(s) declared no potential conflicts of interest with respect to the research, authorship, and/or publication of this article.

## Data availability

Data involved in this work are available upon request.

## Supplementary material

Supplemental material for this article is available online.

## Notes

### Competing Interest Statement

The authors have declared no competing interest.

