## Supplemental Figure 1 for "Bidirectional crusher gradient method for estimating the labeling efficiency of pseudo-continuous arterial spin labeling MRI in mice"

### Supplementary Information

#### 1. Longitudinal contribution of the internal carotid arteries (ICAs) to total blood flow in C57BL/6 mice

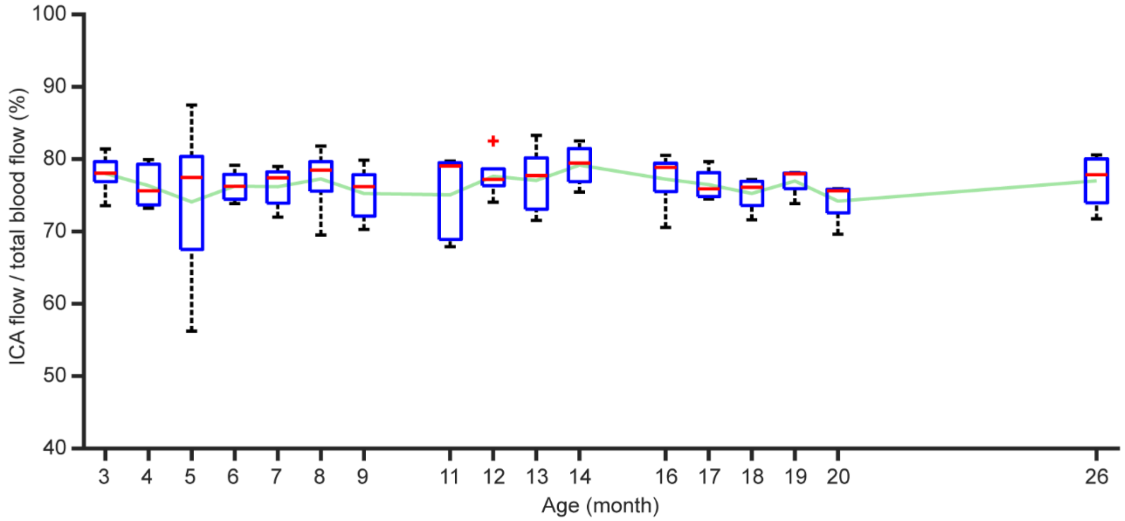


Figure S1 Ratios of blood flow from the ICAs to total blood flow across the lifespan (3-26 month) in C57BL/6 mice (N = 5). In the box plot, the central red mark represents the median, the top and down edges of the box indicate the 25th and 75th percentiles, and the whiskers extend to the minimal and maximal data points not considered outliers. The light green line represents the averaged ratios across 3-26 months of age.

The linear mixed-effect model revealed that the contribution of the ICAs to total blood flow did not show a significant age effect (estimate = 0.061%/month, 95% CI = [-0.070, 0.192], P = 0.355). The averaged ratio of ICA flow to total blood flow was 76.43 ± 1.35% across the 17 time points.
